# The unreachable genomic profiling of complex diseases: genotype missingness matters

**DOI:** 10.1101/2025.07.27.667026

**Authors:** María M. Abad-Grau

## Abstract

The problem of building genome-wide predictors of individual risk to complex diseases seems to be more challenging than it was thought when the first human genome was sequenced on 2003. We have build different enhanced genetic risk predictors from genome-wide data and different complex diseases, making use of haplotypes accurately ascertained from family trios. We confirmed the widely known inability to accurately predict individual risk to complex diseases returned by the state-of-the-art genome-wide predictors. This result is mainly due to the small effect that most genetic variants have in a disease. We also found out that rates of missing genotypes were usually too high for these small-effect variants, as we could force missing imputation in such a tricky way that we would build highly accurate predictors, either by using our own design or the state-of-the-art genetic predictors. We observed that unknown genotypes were not missing at random but related to disease affectation, with more missing genotypes in affected than in non-affected individuals. We were not able to find a way to accurately reduce missing rates to correctly improve accuracy, but we identified a common pattern of missing data in multiple sclerosis, asthma and autism that makes us think that other complex diseases could behave the same way. Because (1) missing rates are high enough to completely change risk prediction due to the small-effect of most of the variants, and (2) there are more missing genotypes in affected than in non affected individuals, we conclude that perhaps the widely known defeat in genomic profiling for complex diseases may be solved by looking closer to the way current genotyping technologies handle genetic variants that may be rare in reference panels but have some effect in a given complex disease.

## 1. Introduction

Since genome-wide association studies started, building genetic predictive models of individual risk to highly polygenetic diseases has became a very challenging task. As an example, in Multiple Sclerosis (MS) the state-of-the-art predictive Area Under the ROC Curve (*AUC*), around 0.64 [1], was achieved by using the common weighted Genetic Risk Score (*wGRS*), while it should be around 0.95 by considering risk prevalence and disease heritability. In fact we theoretically inferred AUC_max_ for MS in Europe to be between 0.9463 (Spain) to 0.9589 (Sweden). We inferred these values from Equations 1 (page 2) and 3 (page 4) in a work publishing the expected AUC for some diseases [2]. We considered sibling recurrence risk λ_S_ = 7.1 [3] and disease prevalence from the European countries with the highest differences: 0.015 in Spain [4] and 0.189 in Sweden [5].

Optimistic opinions about genomic profiling have often flourished to dry soon under the evidence that only under simulations the individual risk can be predicted as expected given the heritability and prevalence of the disease under study.

We began this work after realizing the limits of the state-of-the-art approaches for genomic profiling, such as wGRS [1]. We worked under two generalizations of those common approaches: (1) to fix no limit in the number of SNP loci used in order to take into account very small effects and (2) to use haplotypes instead of just genotypes as a way to keep cosegregation information that may help to better detect risk loci [6,7].

By following these rules, we designed a new genome-wide based method to predict individual risk to a complex disease [8]. We built a very accurate predictor of MS but we were not able to replicate our results in other MS data sets, much less in other complex diseases we tried. At some point after years of tries and defeats, we realized that our first MS predictor was trickily built and our results were wrong. After a few more years of tries, we had to admit that the accuracy of our risk predictors, in agreement with those returned by the state-of-the-art genetic predictors of complex diseases, were far lower than what it should be expected given heritability and disease prevalence of the diseases used.

From so many failures we, by pure chance, discovered an informative missing pattern in all the data sets analysed except for the one corresponding to the Attention Deficit Hyperactivity Disorder (ADHD). All the other diseases analysed during this work have at least 45-50% heritability (twin studies in adults: MS 50-64% [9,10], asthma 75-90% [11,12], autism 50-90% [13] and caries 45-67% [14]). However, ADHD has the highest variability and most discussed in the estimation of heritability, with the lower bound estimation being only around 30% (twin studies in adults: 30-80% [15]).

The causes of the informative missing pattern found in most the data sets analysed is therefore a new research line that has been opened as a result of this long research. In this work we will show the methods used and the experiments performed that have helped to find out this results. We believe this result may be clue to shred some light in the highly challenging task of building genetic profiling for complex diseases from genome-wide data.

## 2. Materials and Methods

### 2.1. Data sets of family trios, genotype calls and quality control procedures

#### Data sets from external Genome Wide Association Studies (GWAS)

We used *IMSGC*, a data set generously provided by one of the authors of a MS GWAS performed on family trios by the International MS Genetics Consortium (IMSGC) [16] with the Thermo Fisher Scientific array (by that time it was Affymetrix) Affymetrix500K. The data set contains the genotype calls and pedigree data of 931 trios once quality control (QC) was performed. There was an affected parent (a mother) in this data set. In a posterior step we also used the IMSGC data set at dbGaP in order to perform genotype calls and to try different parameters for the QC procedures, such as detection of Mendelian inconsistencies and Hardy-Weinberg Equilibrium (HWE), by ourselves. To process raw intensity files, we used the apt-probeset-genotype software provided by Affymetrix with the BRLMM algorithm recommended for its Affymetrix500K array [17]). We also used another data set, *Asthma*, with the genotype calls and pedigree data of the 1323 family trios (after passing QC) from a GWAS of asthma performed by the Genes-environments & Admixture in Latino Americans (GALA I) study, which was also provided by the authors of that study [18]. We also got access through the data base of Genotypes and Phenotypes (dbGaP) to other three data sets with the genotypes and pedigrees of 896, 1323, 178 family trios of Attention Deficit and Hyperactivity Disorder (*ADHD*), autism (*Autism*) and dental caries (*Caries*) respectively. For all of them we used as well genotype calls after passing QC. As the GWAS conducted for dental caries used also case/control subjects and couples we first selected only the 178 family trios whose pedigrees could be clearly ascertained by us. Table 1 shows information about the microarrays used in all these studies.

**Table 1.** Data sets used with sample size after passing QC, genotyping arrays and number of SNPs.

| Trait | # trios | Source | platform | Number of SNPs |
| --- | --- | --- | --- | --- |
| ADHD | 896 | Perlegen | PERLEGEN - 600K | 599171 |
| Asthma | 489 | Affymetrix | AFFY_6.0 | 934940 |
| Autism | 1323 | Illumina | ILLUMINA Human 1M | 1069796 |
| Caries | 178 | Illumina | Human610 Quadv1 B | 601273 |
| IMSGC | 931 | Affymetrix | Affymetrix500K | 262264 (Mapping250K Nsp)<br>238304 (Mapping250K Sty) |
| MS-ES | 80 | Affymetrix | AFFY 6.0 | 934940 |

#### A replication data set for MS from our own GWAS

With the intention to replicate our primary AUC results obtained with the IMSGC data set [8], we conducted a new GWAS from 80 Spanish family trios with offspring affected by MS (*MS-ES*). The Spanish cohort consisted of 80 trios, all natives of Spain. MS patients were diagnosed according to published criteria [19,20]. High-molecular-weight DNA was isolated from whole blood using the Flexigene Kit (Qiagen, Hildren, Gemany) according to the manufacturer’s protocol. Genotyping was performed using Affymetrix 6.0 array at the Spanish National Genotyping Center’s facilities at Santiago de Compostela University (http://www.cegen.org). To keep the conditions as similar as possible, and avoid performing genotype imputation, we used Affy 6.0, the most similar array but that time to the one used by the IMSGC, having a larger set of SNPs but including all the SNPs in the IMSGC data set. We used the Birdseed algorithm provided by apt-probeset-genotype to perform genotype calls in Affy 6.0, the most similar to the one provided by its Affymetrix500K array, BRLMM). We also tried different configurations of the QC procedures, as we did with the IMSGC data set.

### 2.2. Methods

#### Study design

The most widely-used approach to build genome-wide predictors is based on the use of genotypes and the wGRS from which a simple logistic regression model is defined by a given individual *x*: *ln O(x) = ln (p(D|x) / (1 – p(D|x))) = α _0 +_ α _1_ wGRS(x),* with *wGRS(x) = Σ _i_ w _I_ x _i_*, *x _i_* being the individual genotype (0/1/2) at each locus *i, w _i_* the odds ratio at that locus and *O(x)* the odds of the risk of the genome-wide individual genotype *x*. Each individual genotype at a given locus is composed by the nucleotide the individual has at that locus in each homologous chromosome. As the genotyping arrays select relatively-common polimophisms at nucleotide level (Single Nucleotide Polymorphisms or SNPs), for each position there will be only two different nucleotides and therefore the genotype for each locus may be represented by 0/1/2 for homozygotic wild, heterozygotic and homozygotic mutant genotypes respectively. As it can be observer from the definition of the wGRS, it assumes an additive genetic model, as the value *x _i_* of a homozygous individual for the risk allele to the disease at a locus *i,* is double than a heterozygous one.

We used family trio data sets instead of case control ones for two reasons, the first one was to avoid spurious association due to population stratification. The second reason was to use genome-wide haplotypes instead of just genotypes to keep information of real genetic transmission and transcription patterns. Therefore, we defined an haplotype-based wGRS (*hwGRS*), an extension of the wGRS approach based on haplotypes [8]. As we needed highly accurate haplotypes, we had to recover them from family trios.

#### Naive Bayes classifier (NBC) and wGRS

In order to learn the coefficients α _0_ and α _1_ of a wGRS, we assumed that loci are conditionally independent given the disease outcome (D), the assumption made by NBC, and that the two alleles A and B at each locus *i* are identically distributed and conditionally independent given D. Under these assumptions, we probed that

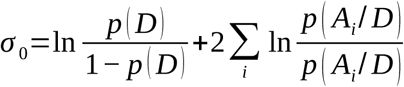

and *σ* _1_=1holds [8]. Simpler coefficient values have been assumed by others [1]: *σ* _0_=0 (no intercept) and *σ* _1_=1.

#### Haplotype-based predictors

We actually built risk predictors using hwGRS for a given haplotype *h*, from which a simple logistic regression model is defined by a given genome-wide haplotype *h*:

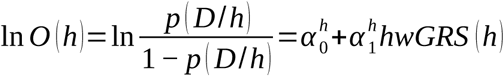

with

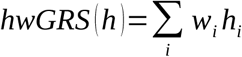

being the allele (0/1) at each locus *i* and *w_i_* the odds ratio at that locus. With the same assumption we did for wGRS, the coefficient values were probed to be:

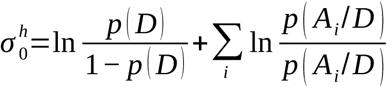

and *α^h^*_1_ =1 [8]. hwGRS is used to independently estimate the risk for the two genome-wide haplotypes an individual has, one from the genome inherited from the father and the other one from the mother. They were combined assuming a recessive genetic model. We also tried dominant and additive genetic models [8]. To graphically understand the difference between hwGRS and wGRS, see Figures 1 and 2 for a wGRS and our generalization, hwGRS, respectively.

**Figure 1.**
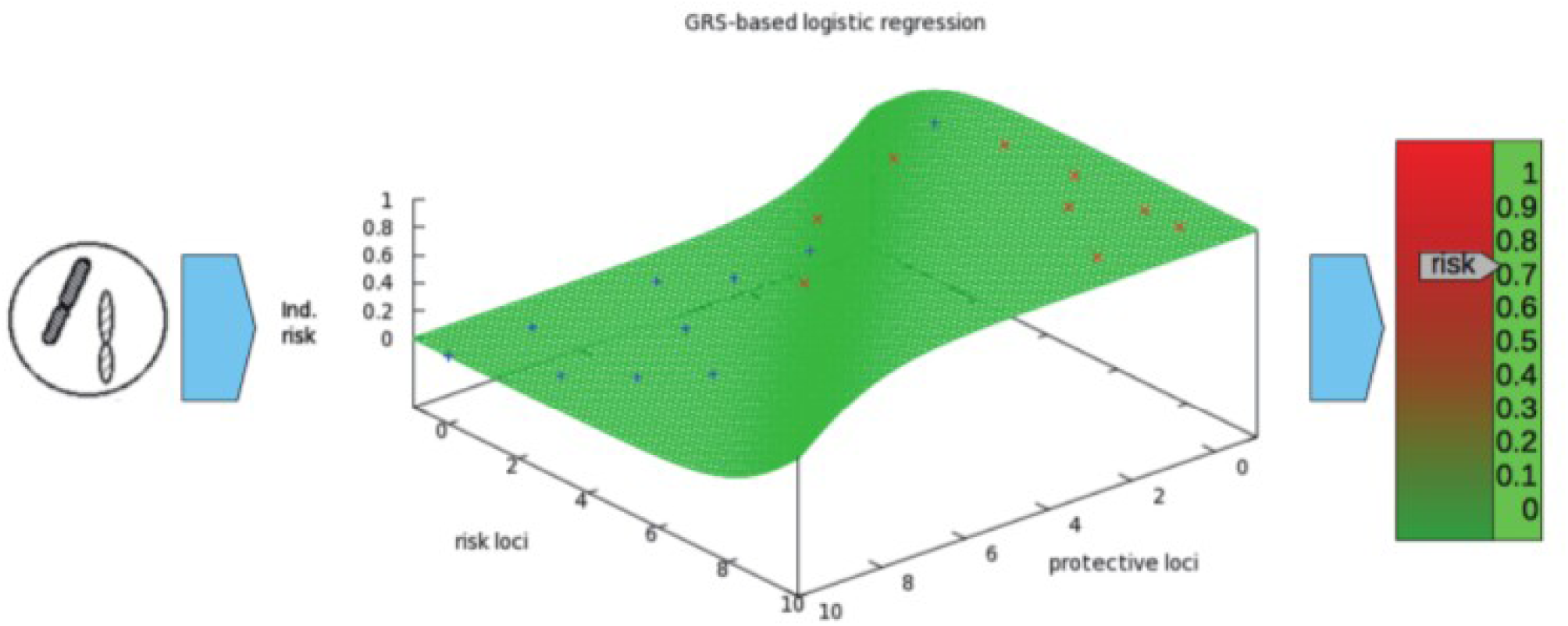
A weighted Genetic Risk Score.

**Figure 2.**
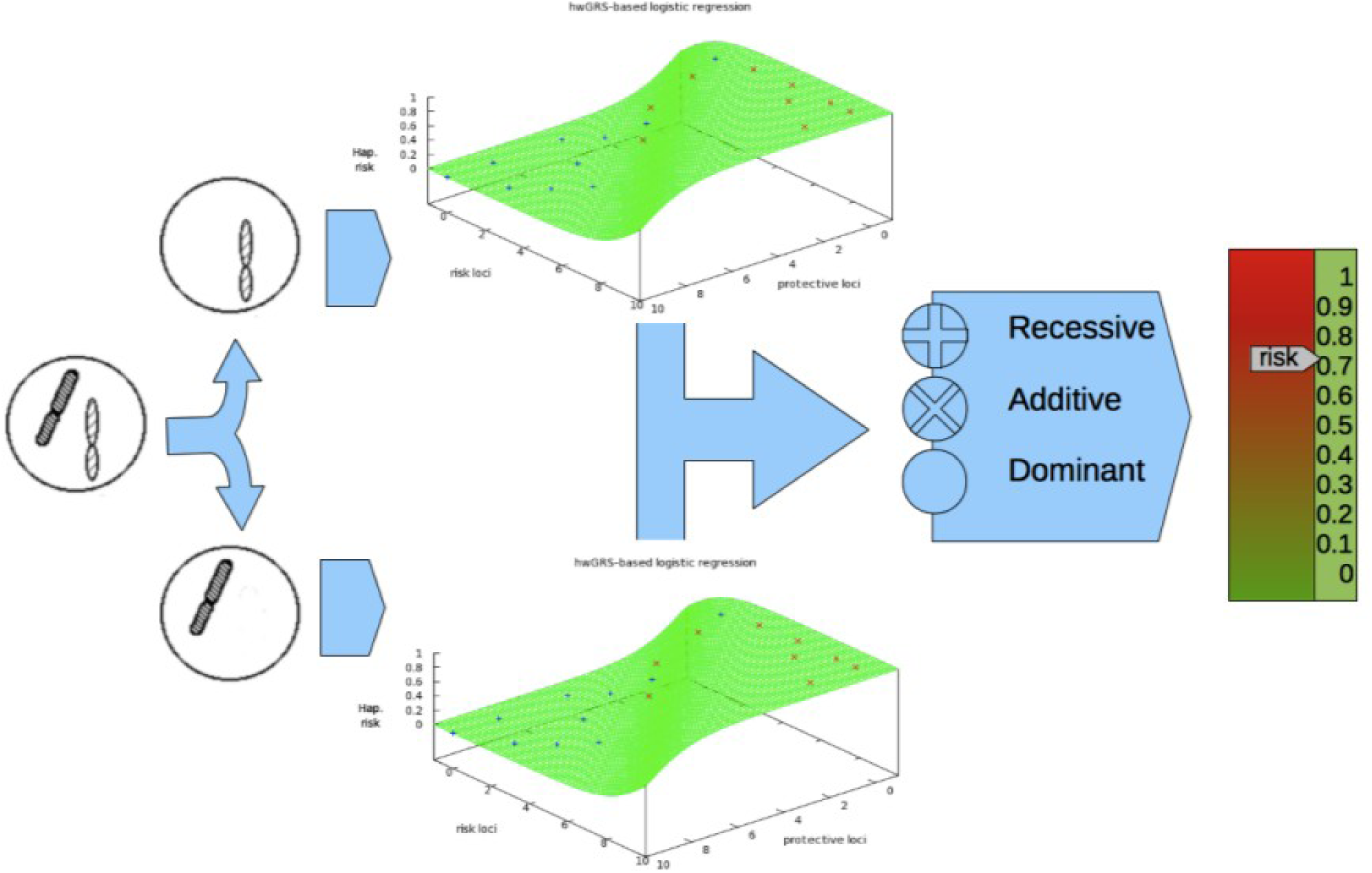
A haplotype-based weighted Genetic Risk Score.

#### Association test

To measure the association between each locus and the disease, we applied *2GTDT* [21], a generalization of the TDT test for family trios, in which association loci has to be detected in two independent subsets of family trios to get rid of spurious association due to random genotyping errors. Therefore, only when it is confirmed in the second subset, the locus was chosen in order to build the predictor. 2GTDT is a multimarker transmission disequilibrium test based on the *2G* approach [21] that is able to collapse the haplotypes in only two groups: high and low risk. By using 2G, the haplotype for each locus in the hwGRS does not need to be a nucleotide but just a binary value resulting from the summarization of haplotypes of any length to only 2 values representing low (0) and high (1) risk of that haplotype to the disease. As it occurs for hwGRS, 2G can also be used as a way to generalize a common wGRS in order to collapse SNPs that are suspected to be in strong linkage disequilibrium (LD).

#### wGRS from hwGRS

We derived the relationship between wGRS and hwGRS from the relationship between the logistic regression for genotypes (wGRS) as the sum of the logistic regressions for each genome-wide haplotype (hwGRS for h1 and h2)) and the *ln* of the odds on the disease:

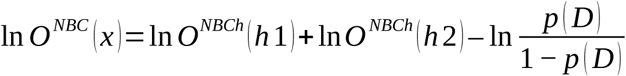

with NBC and NBCh meaning the assumptions made are those of a NBC for three-value (genotypes) and binary (haplotypes) variables respectively [8].

Therefore, instead of using the same 2G approach as a preprocessing step to obtain genotypes representing more than one SNP position, and to compute wGRS afterwards, we can make use of this relationship in order to use 2G as a common preprocessing step for both predictive approaches, if we want to compare outcomes of predictors under the wGRS and the hwGRS approaches.

#### Genetic risk scores and training-test sampling

We built risk predictors by learning the odds ratio of the hwGRS from a training data set and measured accuracy, AUC, sensitivity and specificity on an independent test data set (generalization capacity). Therefore, only a subset of the original data set is used to learn the model. For the IMSGC data set, 500 trios were randomly selected as the training data subsets and the remaining ones made up the test data subset. The number of the randomly chosen trios to make up the training data subsets from the other data sets was 500, 300, 500, and 140 for ADHD, Asthma, Autism and Caries respectively.

#### Sliding windows

In order to better identify significant associations when the causal loci are not in the array but SNPs in linkage disequilibrium with them are, we extended wGRS by using more than one SNP for each input variable. To this end, we used sliding windows of different sizes: 1, 2, 5, 10, 20, 30, 40 and 50.

#### Significance threshold

We used different thresholds of p values to compare the predictor performance. As we used a training-test approach, large thresholds, or even to use all the loci should not reduce the predictor specificity and its specificity could arise as those loci with low or very low effect on the disease are not disregarded.

#### Genome-wide genotype imputation and meta-analysis method

As we were strongly focused on the task of building accurate predictors, close to the AUC expected due to population prevalence and inheritance of each complex disease, we had to suspect of every single state-of-the-art pre-processing step conducted by studies building individual risk predictors (genetic profiles) from genome-wide data. Therefore, we decided to avoid genotype imputation in all the studies conducted, as it assumes that Mendelian errors or departures from HWE are random and this may not be true and thus to provoke crucial information. In order to do that, we try two approaches: (1) to use a data set without passing QC and (2) to follow a meta-analysis method so that QC was independently applied on the original (IMSGC) and the validation (MS-ES) data sets and, afterwards, only the common SNPs between the two data sets after passing QC were selected.

As stated above, 2GTDT is robust to random genotyping errors, as the association has to be confirmed on a second and independent subset of samples than the one used to choose the risk loci.

The second approach, the meta-analysis method, although does not use the validation data set to learn the predictor, is not that good as the first one because there are some shared pre-processing information, as those SNPs removed by the QC step are also removed from the primary data set.

#### Completely independent data sets

Even if we do not perform QC at all and rely only in the robustness of 2GTDT to random genetic errors, it is not true that the predictor is built and tested from two completely independent data sets, as they come from the same GWAS and their genotypes have been called at the same time. We therefore also conducted independent genotype calls for the two data subsets of IMSGC so that they are completely independent.

#### Replication analysis

We estimated accuracy and AUC in a different data set used as a second validation: MS-ES. We only used those genotypes included in the IMSGC data set. As explained above, to avoid imputation we used two different approaches.

#### Ethics Statement

All participants required to obtain the genotypes for the MS-ES data set were recruited and blood extracted during routine hospital visits to the MS unit at Hospital Universitario Virgen Macarena in Seville, Spain, after giving written informed consent under Macarena Hospital Institutional Review Board.

## 3. Results

### 3.1. Predictor performance cannot be replicated

We started this work after doing some tries to build a risk predictor for Crohn disease from a reduced set of SNPs (103) genotyped in 129 family trios and being aware of the substantial advantages of using haplotypes instead of only genotypes, as GRS or wGRS do. This data set was used to show how rTDT [22], a robust extension of TDT to handle missing genotypes, were able to detect possible type I and type II errors when missing data were not ignored.

When we had access to the IMSGC data set, we designed a risk predictor (hwGRS) able to make use of the haplotype information. It returned an AUC value of ∼0.8 when using no filtering for the SNPs selected as input variables and sliding windows had at least 30 SNPs length. This AUC was significantly higher to the state-of-the-art AUC for MS from genetic data (∼0.64), reached by a wGRS built using only 16 SNPs highly associated with MS [1]. Our results were much closer to what it should be expected given genetic prevalence and heritability of MS (∼0.9). We showed that, when using filtering or just genotypes instead of haplotypes, results dropped to the standard AUC results, which supported the idea of using genome-wide haplotypes and a recessive model (see Figure 3 and Figure 4 a).

**Figure 3.**
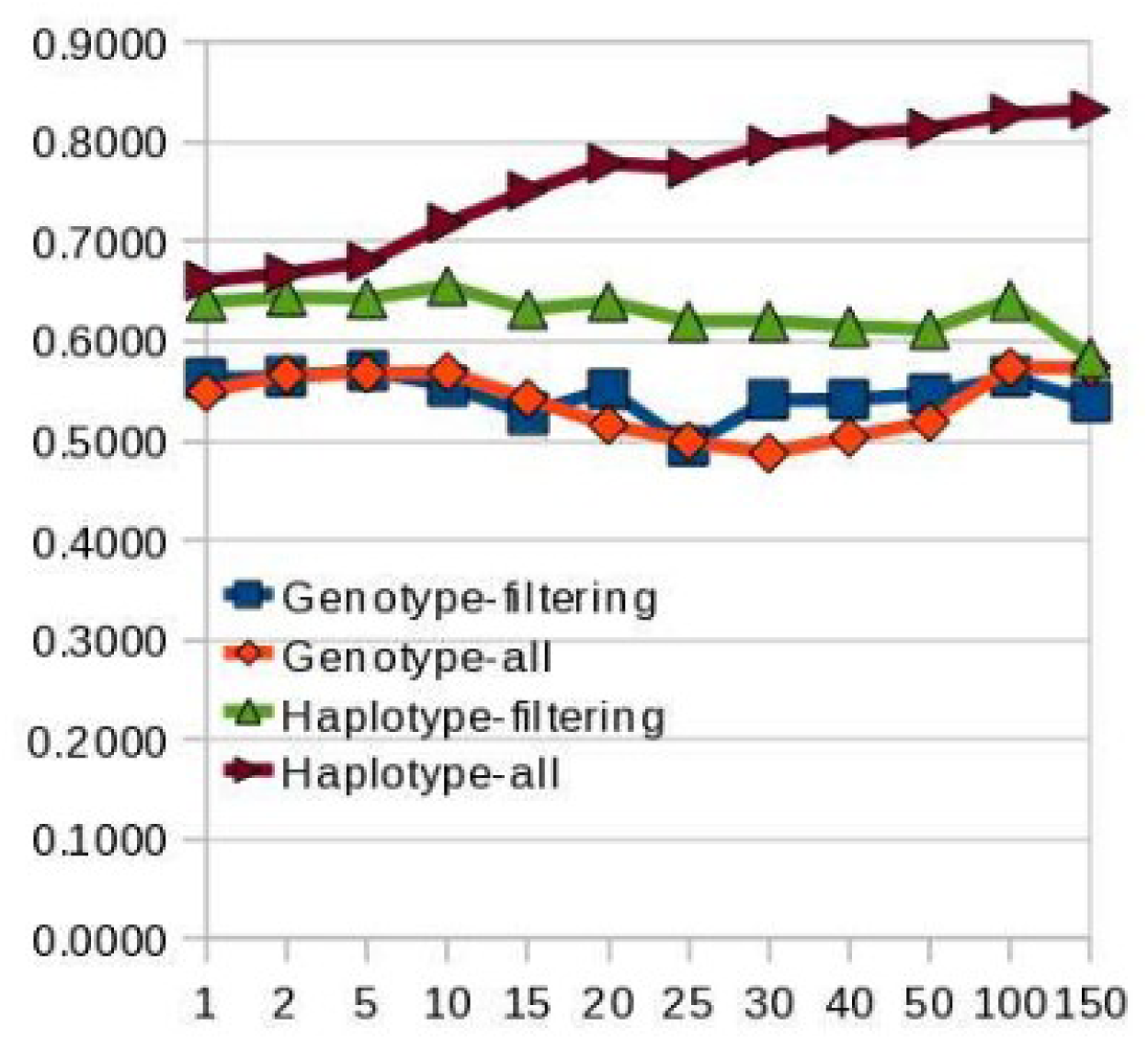
Accuracy (y-axis) of genetic predictors for different sizes of multimarker variables (x-axis). Comparison between hwGRS with no filtering (brown) and filtering by SNPs with strong association to MS (green), and wGRS without filtering (orange) and filtering by the same SNPs (blue). We recently discovered that these results were completely wrong. Source [8].

**Figure 4.**
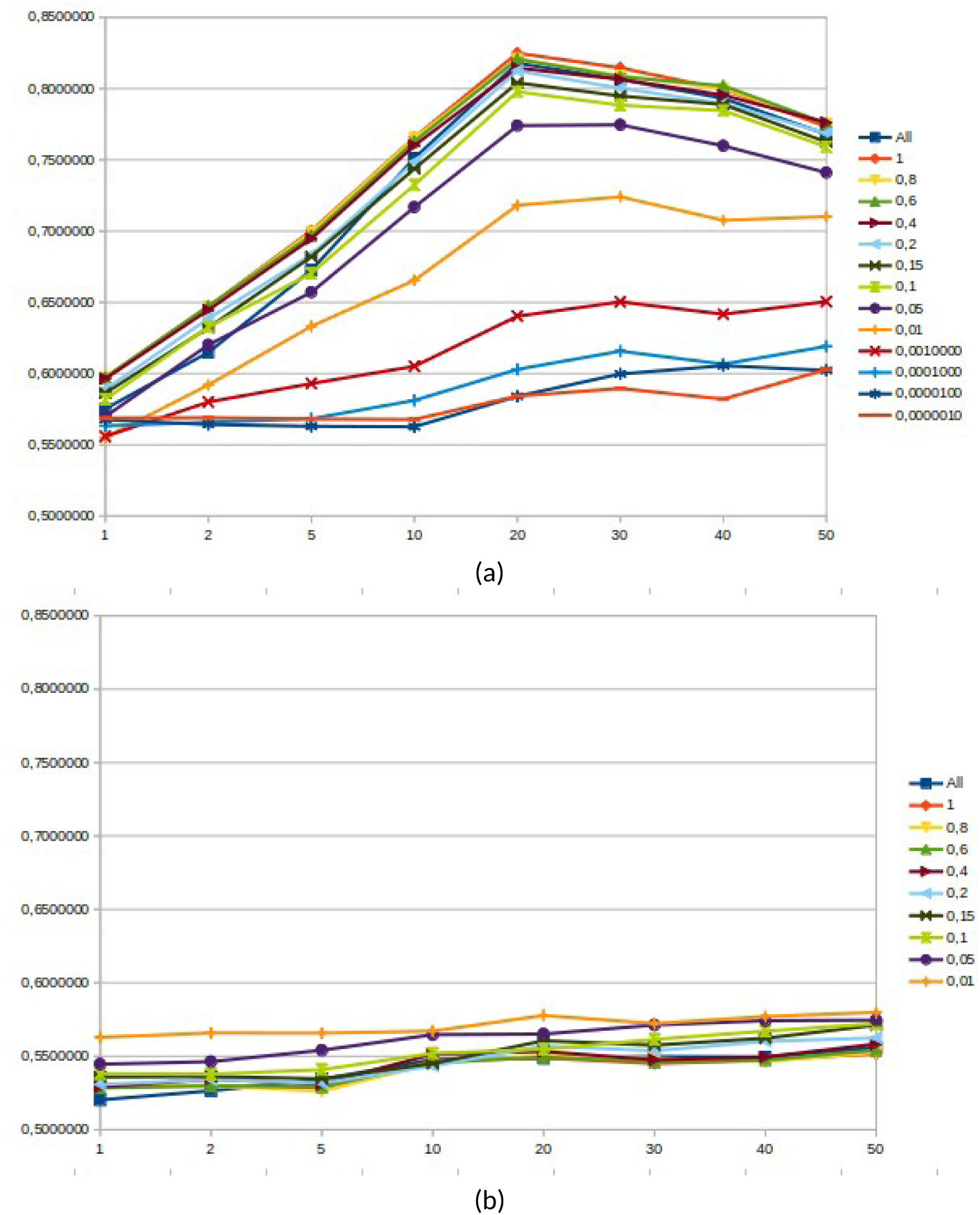
AUC for different p value (from 2G-TDT on the training data set) thresholds (“all” means all SNPs were used) and different haplotype sizes (1 to 50). Model was learned from 500 trios randomly chosen (training data set) and AUC was estimated from the remaining 431 trios (test data set). (a) Results from ped files provided directly by IMSGC before they were released and put in dbGaP. (b) Results from ped files in dbGaP. It has to be noted that source data were almost the same, the different was in the software changes we did to process each data sete by that time, being unaware of them.

As we were highly concerned by this highly astonishing AUC reached by hwGRS under a recessive model, we only dared to publish our method and results as a work in process [8]. We decided to perform our own GWAS (data set MS-ES) with the intention to confirm and replicate this result. We only used 80 trios because of budget limitations.

However, after many tries, we were unable to obtain good results at all (AUC was below 0.65) and for about 2 years were completely unable to understand the reason. What it was worse: when we tried to repeat the experiment with another version of the IMSGC, obtained from dbGaP, we could not replicate the first results (see Figures 3 and 4).

This low generalization capacity was in agreement with results obtained by other state-of-the-art multiplex predictors [1,7] and it is against the theoretical AUC estimated for several highly complex diseases, in which there may be thousand predictors of low or very low disease effect [2].

The cause could not be attributed to an overfitting problem, a problem that arises when the learning algorithm used to learn the model does not take into account a threshold between the sample size of the training set used to build the model and the model complexity itself. In fact, we, as others [7], used a training-test method to avoid overfitting. Moreover, a weighted genetic risk score is equivalent to a naïve Bayesian network [23], the most simple model for a Bayesian network, as it does not consider between-variant dependencies given the disease outcome. If the training data set is too small to capture some small-effect variants, the weight (usually odds ratio) for these variants will be commonly underbiased and sensitivity results measured in the test data set will be very low. This is the main issue that most multiplex predictors of complex diseases have and why a possible solution to improve their low predictive capacity is not to change the algorithms used to learn the multiplex predictor but to significantly increase the number of individuals genotyped in order to learn the model.

At some point we realized the software implementation by the time we started using the other data sets, including the IMSGC data set downloaded from dbGaP, had changed a lot and the most important difference was that the missing data were not imputed in the way we did with the IMSGC data set the first time we built our predictor. We focused therefore in the task of discovering what was the difference between the two versions of IMSGC we had.

We decided to try at least to extend these results to any of all the other diseases for which there was a trio data set that we could get access through the NIH dbGaP: caries, ADHD and autism. We also had a data set of asthma in a Latino population [18]. We were not able to obtain noticeable results for AUC under the same training/test sampling approach. We actually obtained quite low AUC values, by considering all the configurations of window sizes and p value thresholds used for MS (∼ 0.60-0.65). For ADHD the results showed an almost null genetic effect (AUC values were very close to 0.5, which means there is almost random guessing). For technical reasons we could not finish testing the predictors in autism and caries.

On September 1st 2016, we first time were able to understand the way we were imputing missing genotypes in the first version of IMSGC data set. We even enhanced the preprocessing steps and were able even to increase AUC to values closer to that expected for MS by considering disease prevalence and heritability (∼0.90). However, at that time, we thought that it was not a bias but the correct way to impute missing data introduced by us by pure chance and therefore we should do the same with MS-ES, the Spanish data set. We were very confident we would now be able to replicate our results.

It was a step forward, even if we were still really far away from the truth. We were able to impute missing data in almost the same way we did first time. We actually coded a better implementation of what it was done by chance the first time: to impute data in order to favour the transmission of the allele that, in the known genotypes, is the less transmitted one -sometimes just called the minor allele under the TDT approach-, an imputation method we called *cT*. The AUC was significantly higher and we finally were able to obtain the same results from the IMSGC data set version we downloaded from the dbGaP server.

The most important knowledge learned in this step was that missing data was crucial for the correct performance of a genome-wide risk predictor, i.e., to reach an AUC close to the theoretical value. When using the cT approach in the replication dataset (MS-ES) we were able to reach similar AUC rates. When we used the cT approach with ADHD data set and AUC did barely improve. This could support the claims that genetics on ADHD is not that important [15].

### 3.2. Studying the missing pattern

We were very suspicious and afraid of publishing highly positive results that could be wrong. We decided to try first a different experiment to check whether the missing patterns were consistent in both data sets. Because of this, we employed some time trying to understand the missing pattern and we made other some interesting discoveries. We showed that the missing pattern, before any transformation, in the IMSGC (Figure 5) and MS-ES (Figure 6) data sets was not random but informative. In fact, there were more missing genotypes in affected individuals (offspring) than in unaffected (parents) ones in all the chromosomes. As there was one affected parent we showed that missing rates for this individual were, for all chromosomes, closer to average rates in offspring than in the other parents (Figure 7). We also studied the pattern of Mendelian errors and observed they were skewed towards the minor allele in all the chromosomes (see Figure 8).

**Figure 5.**
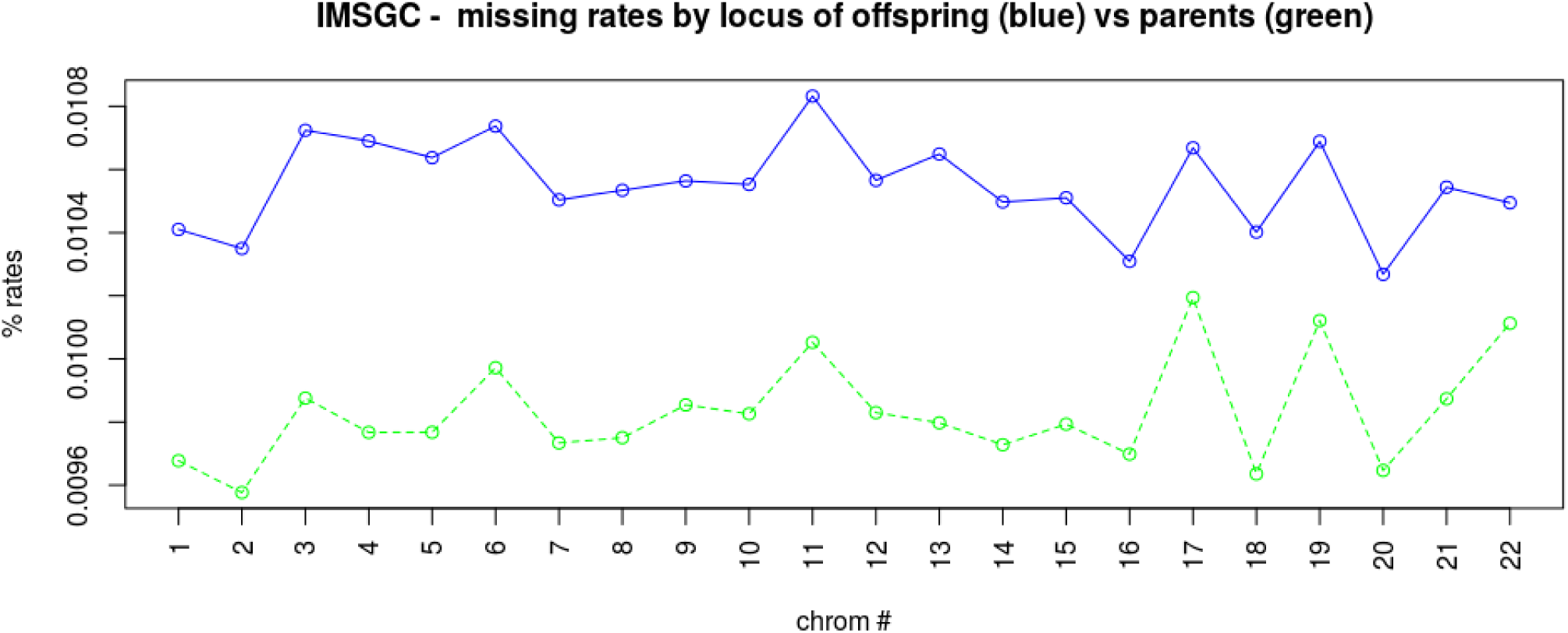
Line plots with average missing rates by chromosomes for parents and offspring in IMSGC MS data set. Plots were produced by using the whole data set with 931 trios.

**Figure 6.**
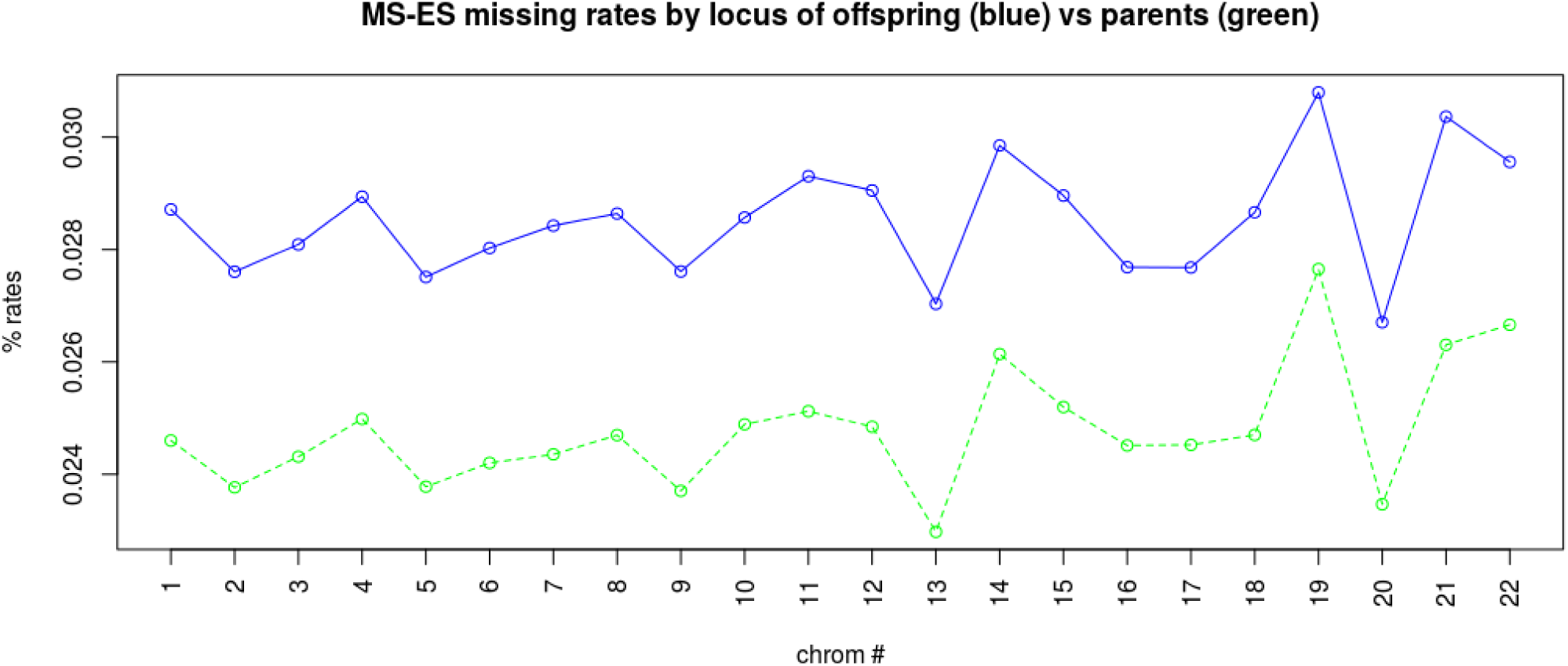
Line plot with average missing rates by chromosomes for parents and offspring in the Spanish MS data set. Plots were produced by using the whole data set with 80 trios.

**Figure 7.**
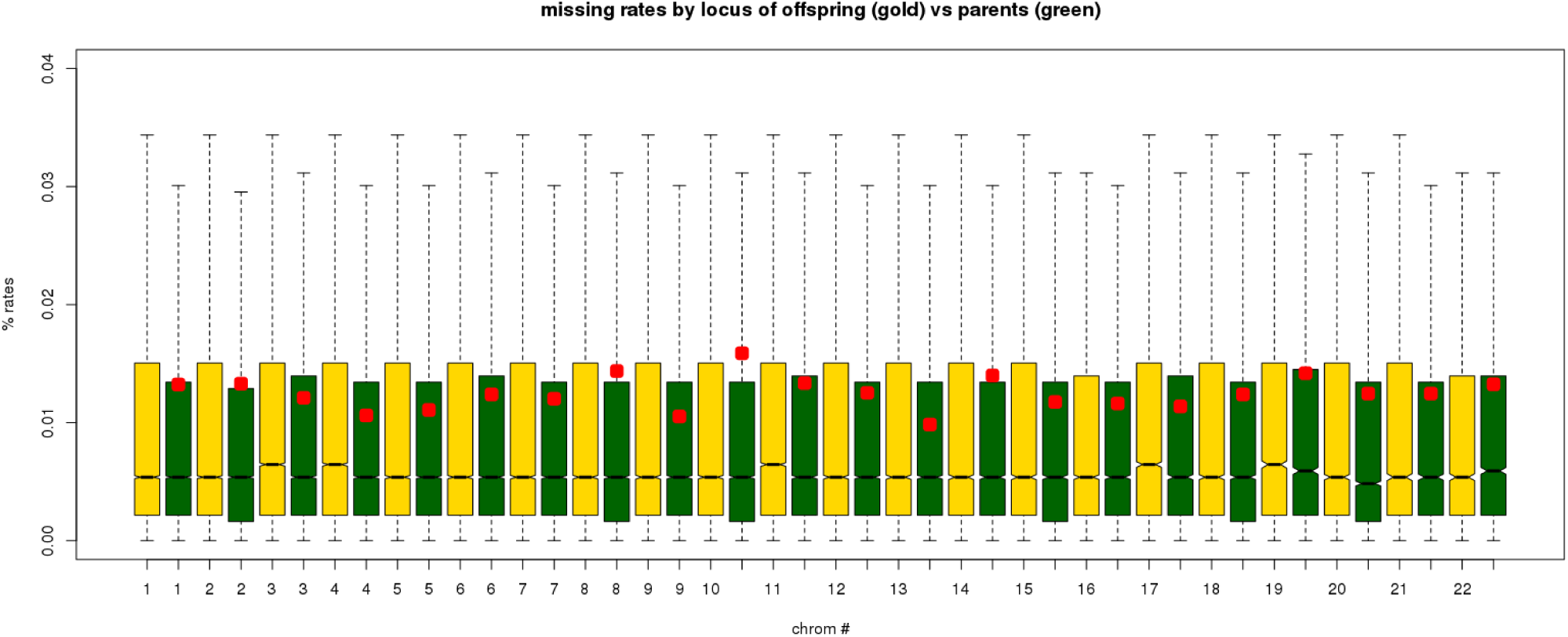
Box plot by chromosomes for parents and offsprings in IMSGC data set. The average missing rates for the affected parent are shown in red.

**Figure 8.**
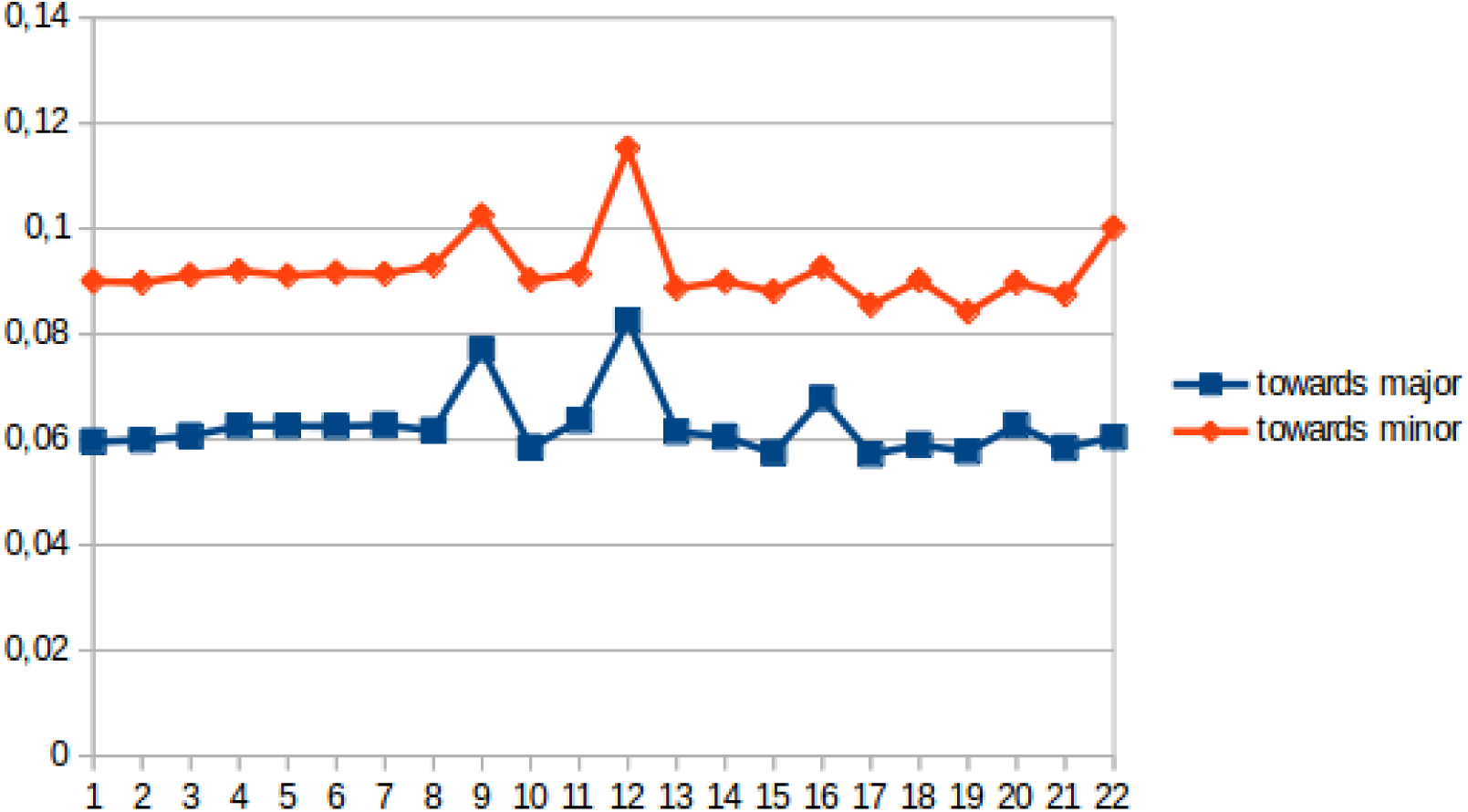
Line plot with average Mendelian error rates by chromosomes towards major (blue color) and minor allele (red color) in IMSGC data set.

We confirmed the same missing pattern found in the two MS data sets in other two data sets we got accessed to: asthma and autism (see Figures 9 and 10 respectively).

**Figure 9.**
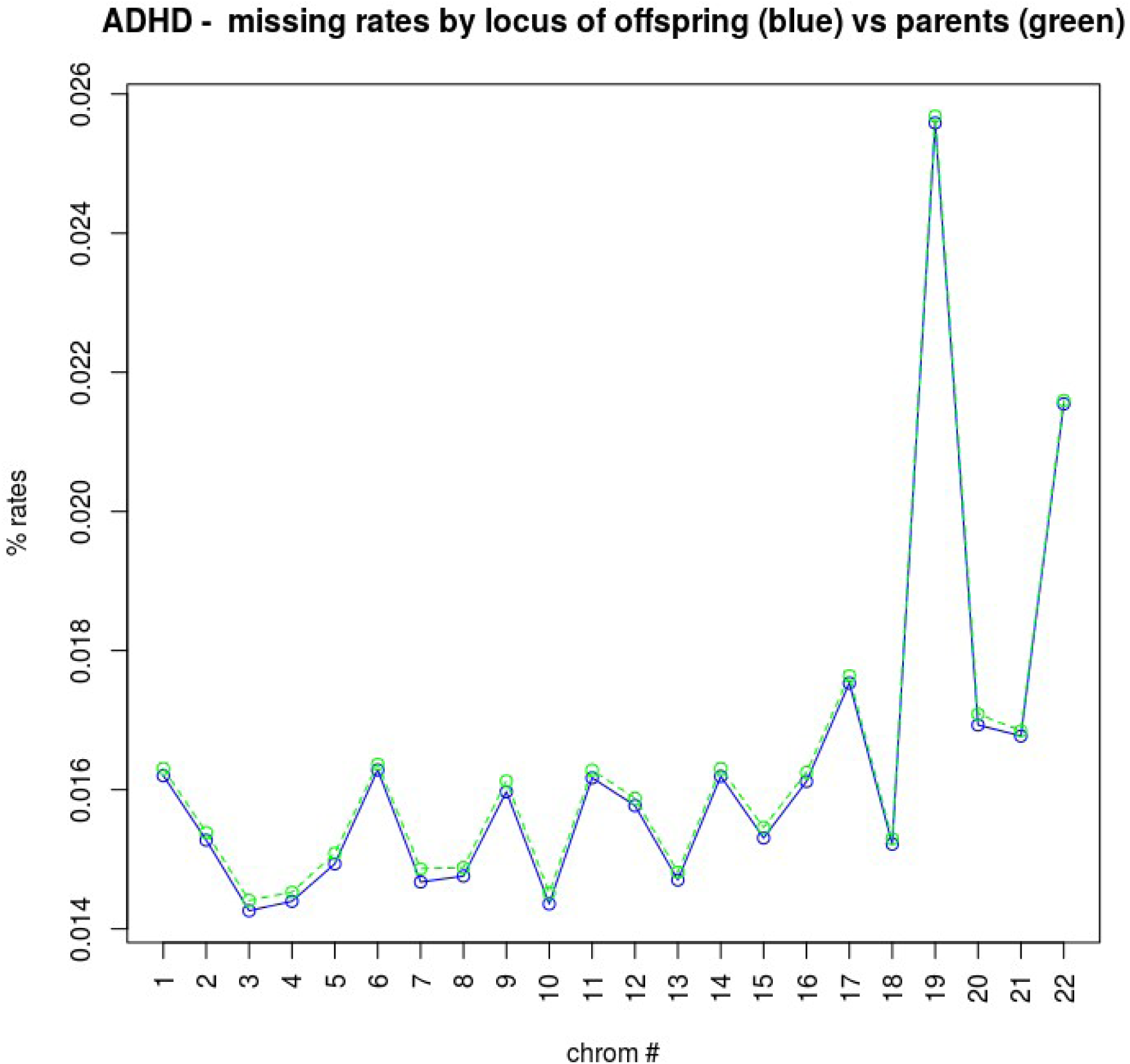
Line plot with average missing rates by chromosomes for parents and offspring in ADHD data set. The plots was produced by using the whole data set with 896 trios.

**Figure 10.**
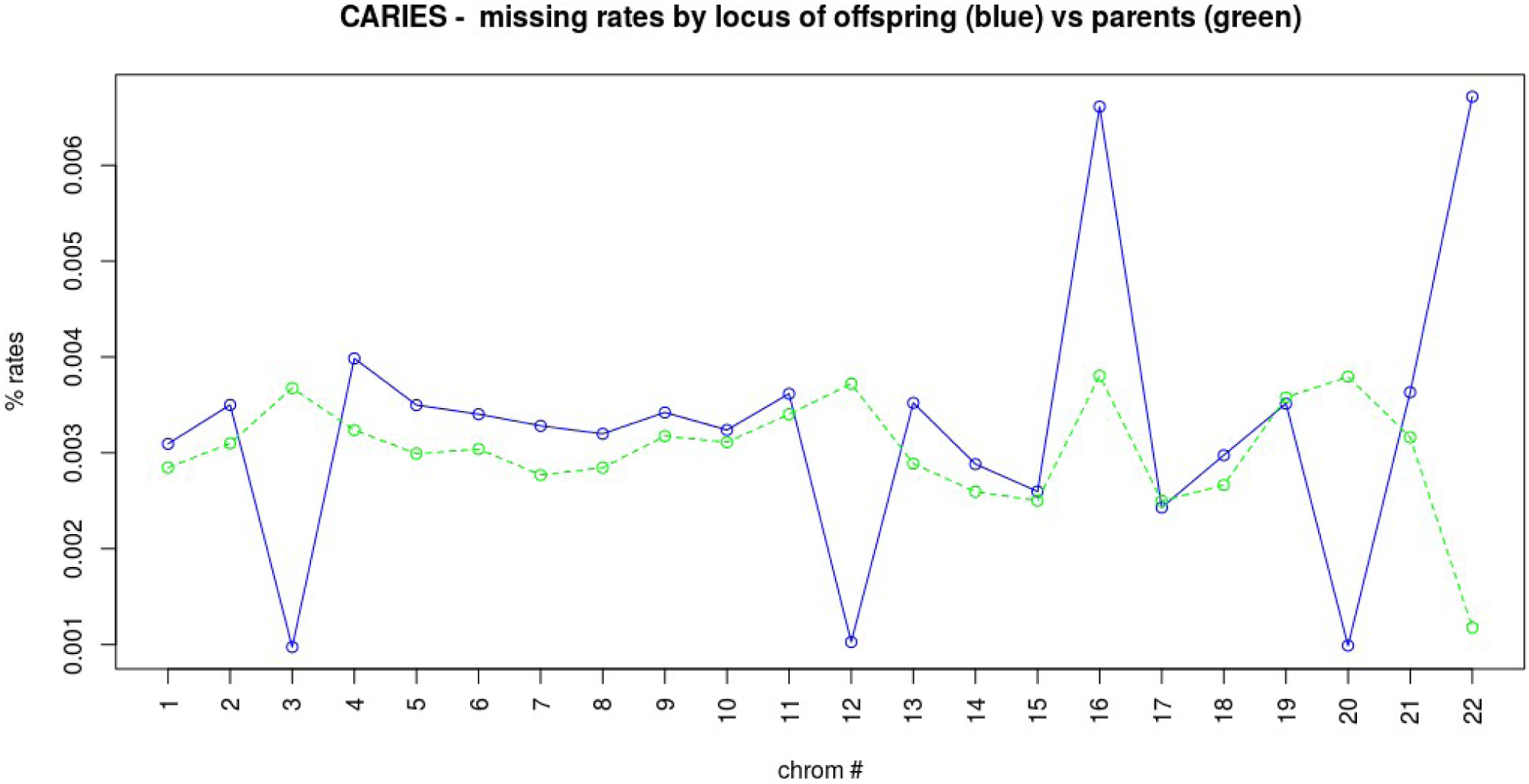
Line plot with average missing rates by chromosomes for parents and offspring in Caries data set. The plots was produced by using the whole data set with 178 trios.

We also observed that the pattern did not hold in ADHD data set (Figure 11), which was consistent with the low AUC on the test data set returned by the predictor we built.

**Figure 11.**
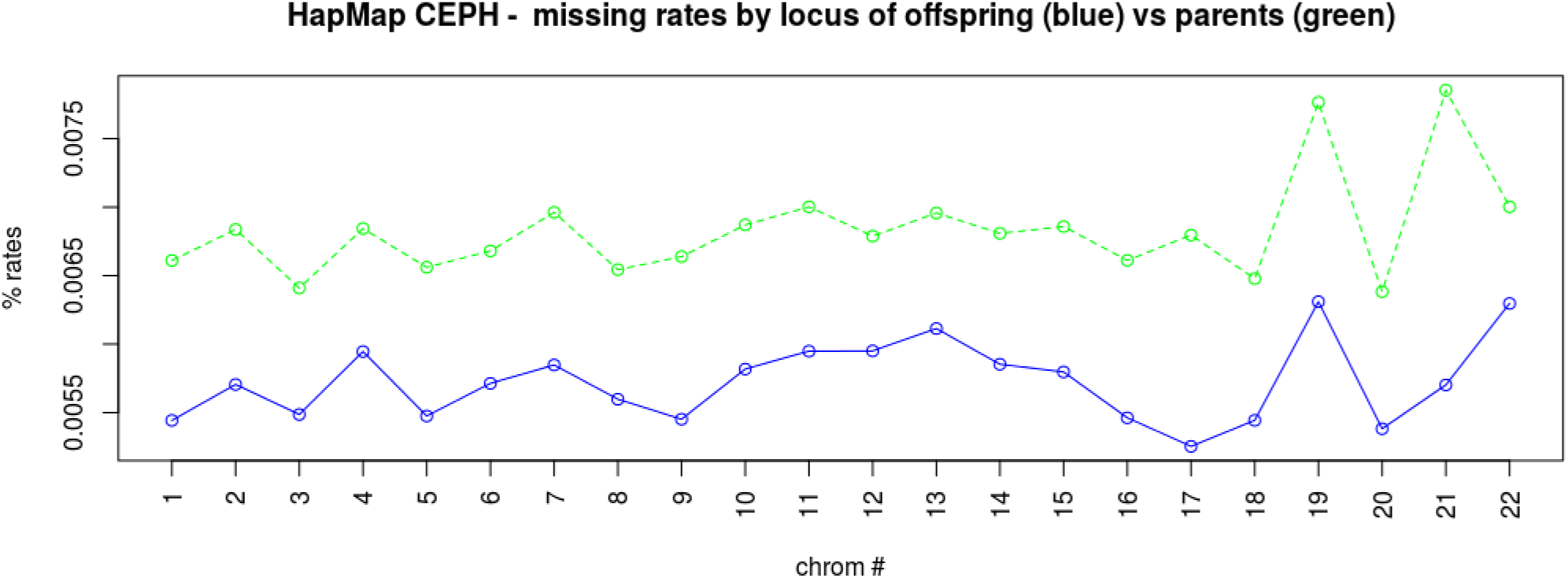
Line plots with average missing rates by chromosomes for parents and offspring in HapMap CEPH data set. Plots were produced by using the whole data set with 30 trios.

When using the caries data set, we observed the pattern was not consistent among chromosomes (see Figure 12). As we did not have performance results for the caries predictor, we did not make any conclusion about it.

**Figure 12.**
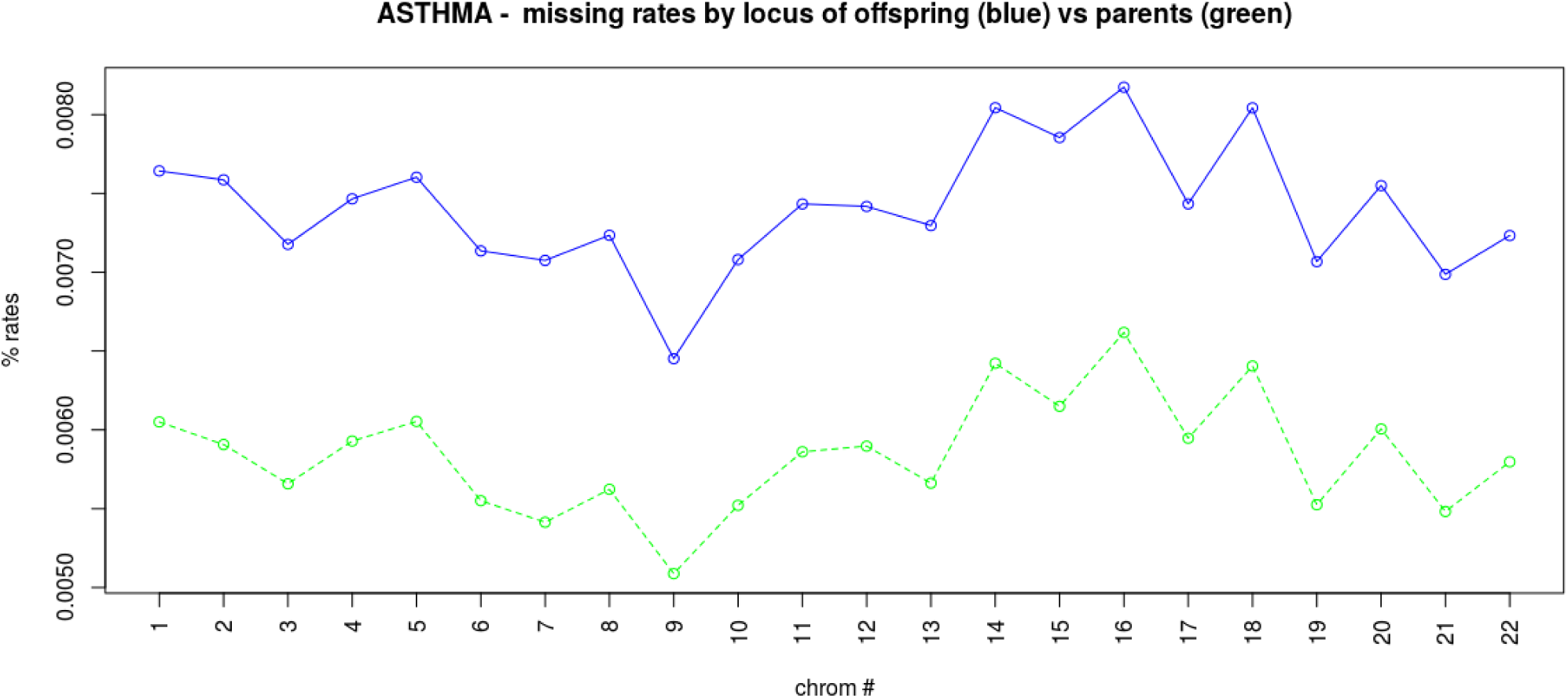
Line plots by chromosomes for parents and offspring in the asthma data set.

In order to check that the informative missing pattern was different in a trio data set of healthy individuals, we also made the same plot for the CEPH data set from the HapMap project (Figure 13). It was very interesting to observe a consistent informative missing pattern among chromosomes but exactly in the opposite way: there were more missing genotypes in parents than in children. The explanation seemed to be clear: in a trio data set, the genotypes of offspring appear twice (once in the offspring, once in their parents -half in the father and the other half in the mother-) while only half of the parental genotypes are repeated (the part transmitted to their offspring). Therefore, their alleles become easier to be ascertained by the genotype calling algorithms.

**Figure 13.**
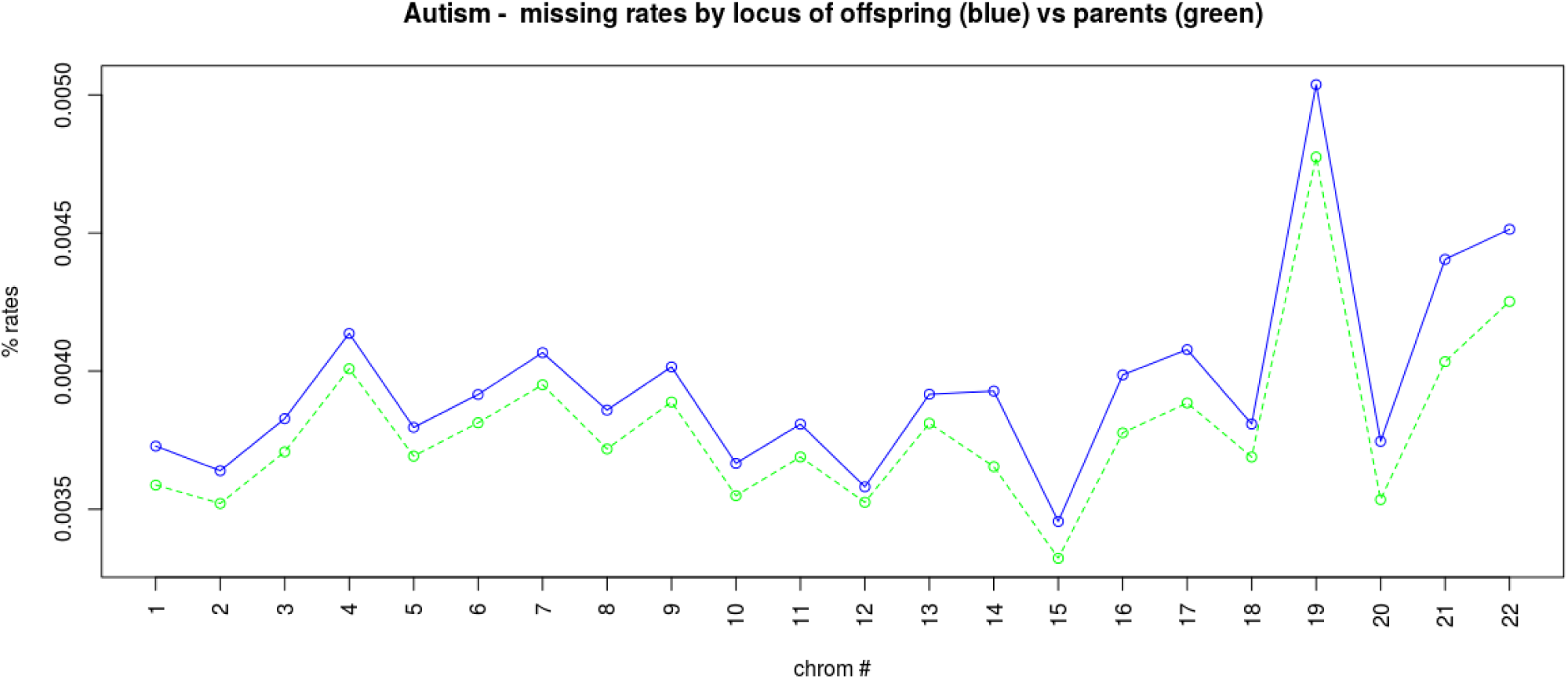
Line plots by chromosomes for parents and offspring in the autism data set.

### 3.3. A highly upbiased predictor: identifying the sources of error

At some point on October 2016 we realized that neither the cT approach or MUT (another approach we tried in which missing are completed in order to increase the transmission of the mutant allele), could not be used for the test data set, as it imputes parents -most of them are unaffected and therefore used to compute specificity-in a different way than offspring.

Moreover, in the way we imputed data, instead of reducing type I errors and make the predictor most robust to missing data even at a cost of a lower sensitivity, we were doing exactly the opposite for SNPs with a low or very low contribution to the disease: increasing type II errors and upbiasing the predictor sensitivity. This was due to a wrong implementation of the cT approach. In fact, our first computer programs for genome-based studies with family trios imputed missing data as their default configuration, as most algorithms did not allow missing values. Trying to avoid type I errors even at a cost of a higher level of type II errors, the missing data were imputed in order to reach the minimum value of rTDT [22], an approach that we have called cT in this work. However our implementation was too simple and actually wrong for genome-wide data sets. By that time and because our only experience was with a short data set of only 129 trios and only 103 SNP (the Crohn data set) [22], we missed some issues to this implementation that would only arise with larger data sets and a greater number of SNPs, as the IMSGC data set used in the work here described. In fact, we imputed missing data at a given SNP locus in order to maximize the transmission of the minor allele. By that time, we did not take into account that, in the case of many missing genotypes, imputing all of them the same way may switch the minor allele into the major allele. Therefore, by computing the TDT statistics *t* this way on these data, rather than being equal to *t_min_* -that it should be actually 0-, it may become much larger than *t* without missing imputation. When we were still unconscious of this big issue of the cT approach and we reimplemented the software trying to better apply the cT approach, the upward bias was actually higher. In fact, undert this approach, for SNPs with a low or very low effect to the disease, *t* is actually estimated almost as the *t_max_* interval upper bound in rTDT, so that we would get much more false positive SNPs associated with the disease.

### 3.4. Reviewing the methods used to build the predictors

On light of this result, we started trying many different ways to perform genotype calls, impute missing data, select training and test data subset, etc. in order to obtain an AUC for MS closer to its expected value considering disease prevalence and heritability.

We tried to keep missing genotypes, or, once alleles are completed from the family trios, to impute the still missing alleles with any method that do not differentiate between parents and offspring. As an example we tried to impute by using the mutant or minor allele (MU approach) or the *c* allele (the one with the lowest transmission). We also tried *MUexp*, an approach consisting on imputing missing genotypes by the minor allele but only genotypes randomly chosen in a proportion given by the exponential function *y=0.000001 exp x*, with *x* being the p value of the test that measured missing differences between affected and unaffected individuals within the training data set. As an example, for a SNP with p value 0.1, the proportion of genotypes to be randomly chosen to be imputed by the minor allele would be ∼25% while for p value 0.05 it would be ∼50%. When we tried to implement this solution we decided to apply it before imputing missing genotypes in offspring that can be inferred from their parents, in order to avoid altering missing rates in offspring before applying the MUExp method because of family data.

We also introduced a change in the way to impute missing genotypes. So far, we had never allowed or introduced Mendelian errors when imputing a genotype. This time we thought about allowing it, as Mendelian errors seemed to be skewed towards the minor allele (Figure 8) and therefore it could be an important cause of individual risk that we should not disregard. In fact, together with this, we decided to enhance TDT in order to measure skewness of Mendelian errors towards one allele. The way to do that is just to “correct” Mendelian errors in the training data subset by changing parental alleles. As an example, if a trio genotype is 11/12/22 (father, mother and offspring respectively) the correction would change the father genotype for it to be heterozygous: 12/12/22. This way, both parents would contribute to increase counts in 2G-TDT.

We tried these approaches and many others without any success at all.

### 3.5. Trying to recover information from the missing genotypes

We decided to have a closer look to the missing pattern so that we could observe if it depends on the known genotypes of the other family members in the trio. Under this criterion, there are only three of them that can be imputed with no ambiguity: those with both parents homozygous (see Figure 14). There are 17 different types of missing patterns with ambiguous missing genotypes (see Table 2 at the work about rTDT [22] and Figure 15). These plots helped to understand that the minor allele (represented by the value 2) was more difficult to ascertain by the algorithms that make the genotype calls. It has to be noted that it was more difficult to make the call for a parent with the minor allele than for an offspring. In fact, missing rates for a trio (0,0) (2,2)(2,2) (Figure 15) is double (∼0.007) than for a child (2,2)(2,2)(0,0) (Figure 14). We cannot be sure whether the missing parent in trio genotype (0,0)(2,2)(2,2) is truly (2,2), as it could be also heterozygous. However, given the high missing rates, it seems that they would be most probably (2,2).

**Figure 14.**
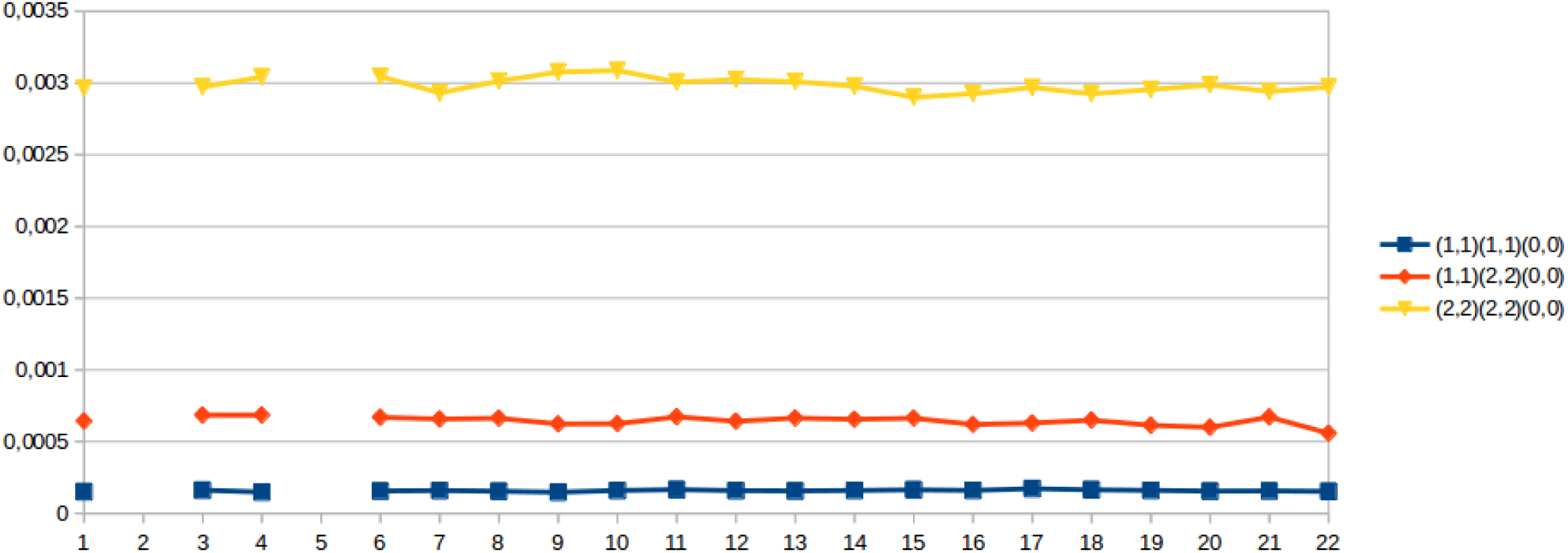
Missing rates in the three missing trio genotypes that can be completed with no ambiguity.

**Figure 15.**
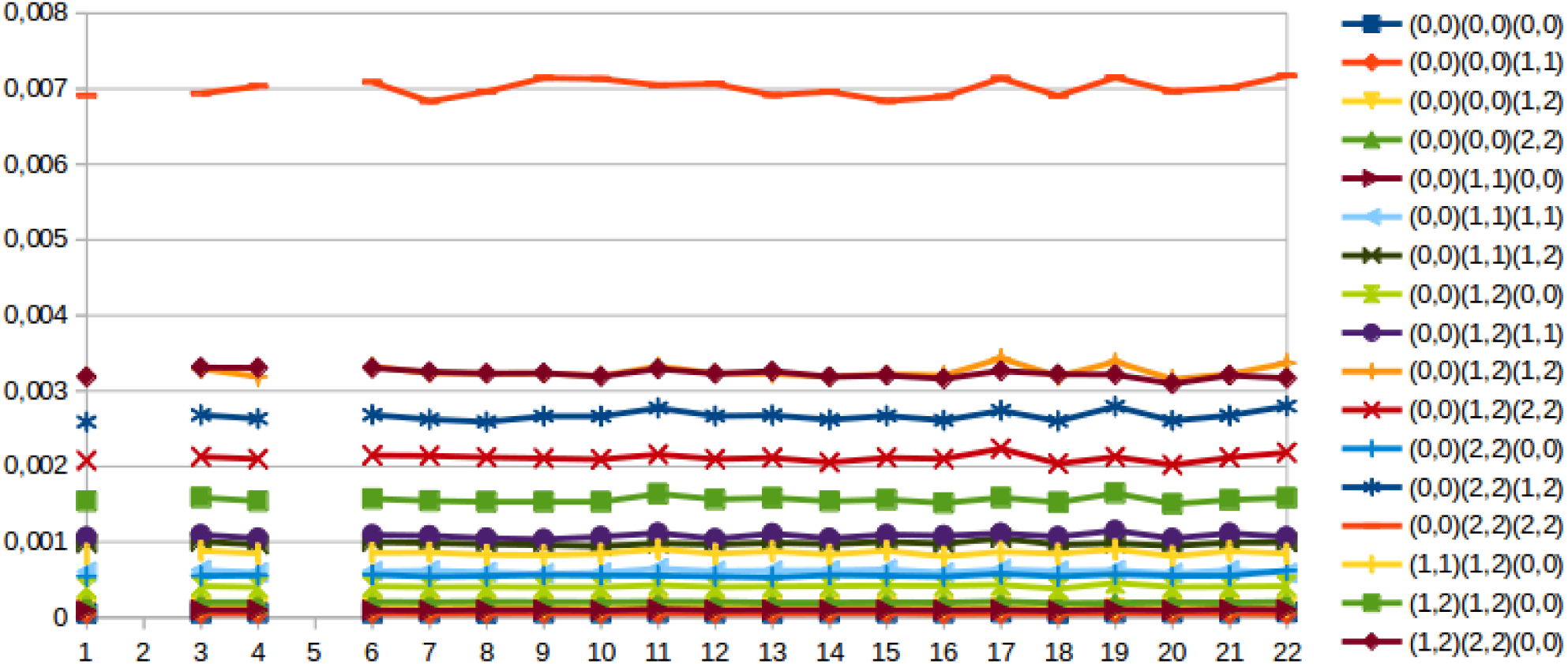
Missing rates in the 17 missing trio genotypes that can be completed in different ways.

We also had a look to the raw intensity values and how genotypes were solved or left as missing by the algorithm that made genotype call. We wanted to observe whether there can be different patterns of missing genotypes. BRLMM [17], the one used by the IMSGC data set, which is based on DM [24]. DM considers there are four models for genotypes: heterozygous (1,2), homozygous wild (1,1) homozygous mutant (2,2) and null (when neither of the other three ones hold). We selected from MS-ES the five SNPs with higher missing rates in offspring than in parents (p vals from 1.129e-05 to 8.707e-05). We discovered that in three out of the five SNPs (see Figure 16, plots corresponding to SNPs SNP_A-8311817, SNP_A-1933659 and SNP_A-2015495), the three-genotypes assumption was far from being observed in a real situation. In order to identify and understand different possible patterns of missing genotypes, we should understand first these patterns of intensities. For a comparative purpose, we also made plots for the first five SNPs with more missing rates in parents than in offspring (as expected, p vals were significantly lower: from 0.0005242 to 0.0022759). As it can be observed (see Figure 17), only one of the SNPs (SNP_A-1851845) does not show the three-genotypes pattern. However, in this case the calling algorithm has not been able to get trustable genotypes for most of the individuals (there are an enormous number of missing genotypes) as the rare patterns affect to many individuals.

**Figure 16.**
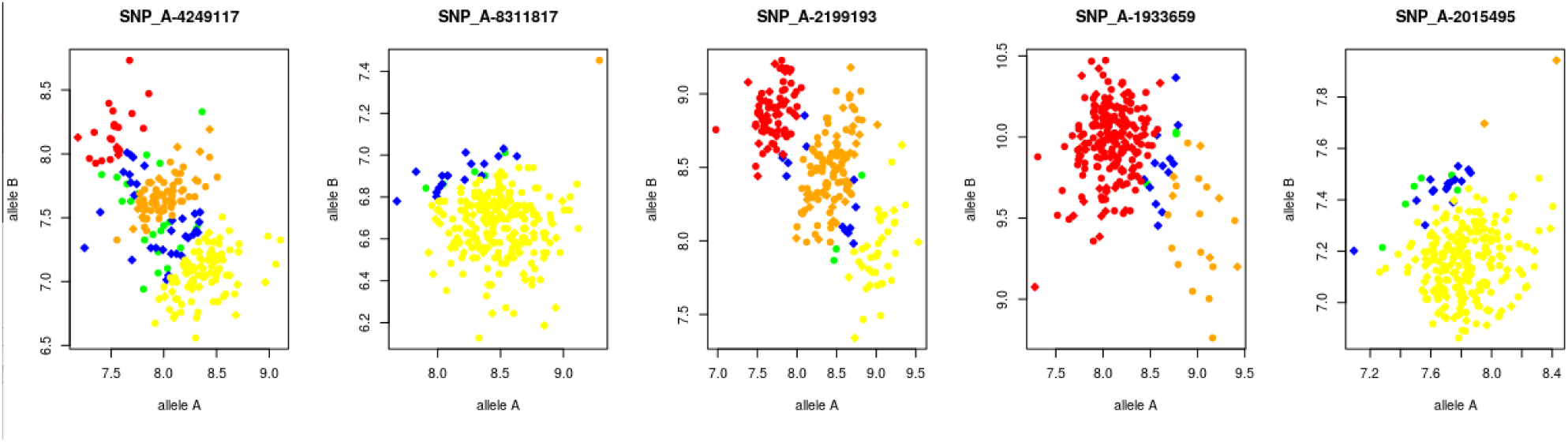
Scatter plots with intensities of A (x-axis) and B (y-axis) alleles in the five SNPs with more missing genotypes in offspring compared to parents in the Spanish MS data set. A green circle means a missing parental genotype while a blue diamond means a missing offspring genotype. Homozygous type 1 genotypes are shown in yellow, heterozygous genotypes are shown in orange and homozygous mutant are shown in red (circles for parents, diamonds for offspring).

**Figure 17.**
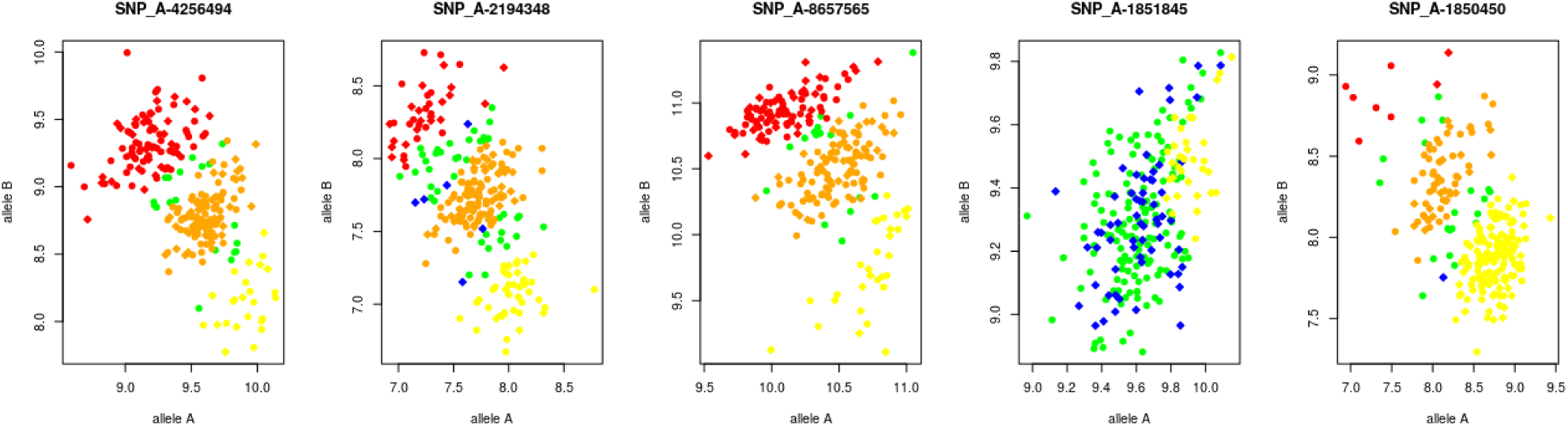
Scatter plots with intensities of A (x-axis) and B (y-axis) alleles in the five SNPs with more missing genotypes in parents compared to offspring in the Spanish MS data set. A green circle means a missing parental genotype while a blue diamond means a missing offspring genotype. Homozygous type 1 genotypes are shown in yellow, heterozygous genotypes are shown in orange and homozygous mutant are shown in red (circles for parents, diamonds for offspring).

### 3.6. Genotype calling algorithms

We looked to the parameters that can be used with apt-probeset-genotype, a free implementation of BRLMM provided by Affymetrix. Among the many parameters that could be used, we saw the possibility of learning a clustering model for each SNP (the three centers and variances) from a data set and to use these models to call genotypes once the allele intensities were obtained. We made use of this possibility in order to understand if missing rates would change. We used only offspring from half of the training data randomly chosen to learn the clustering models. When we used these models to make the calls for the same samples and their parents, the missing rates differences changed the sign, this time with more average missing rates in parents than in offspring (see Figure 18). When we used the same model with the test data, again the common pattern of higher missing rates in offspring in all the chromosomes arised (see Figure 19). The explanation was in agreement with the already known genetic pattern of MS. The fact that using an independent set of offspring to learn the models and make the calls gives higher missing rates in offspring was showing that the variants related to MS risk in one offpsring data set may not be the same than those in the other offspring data set.

**Figure 18.**
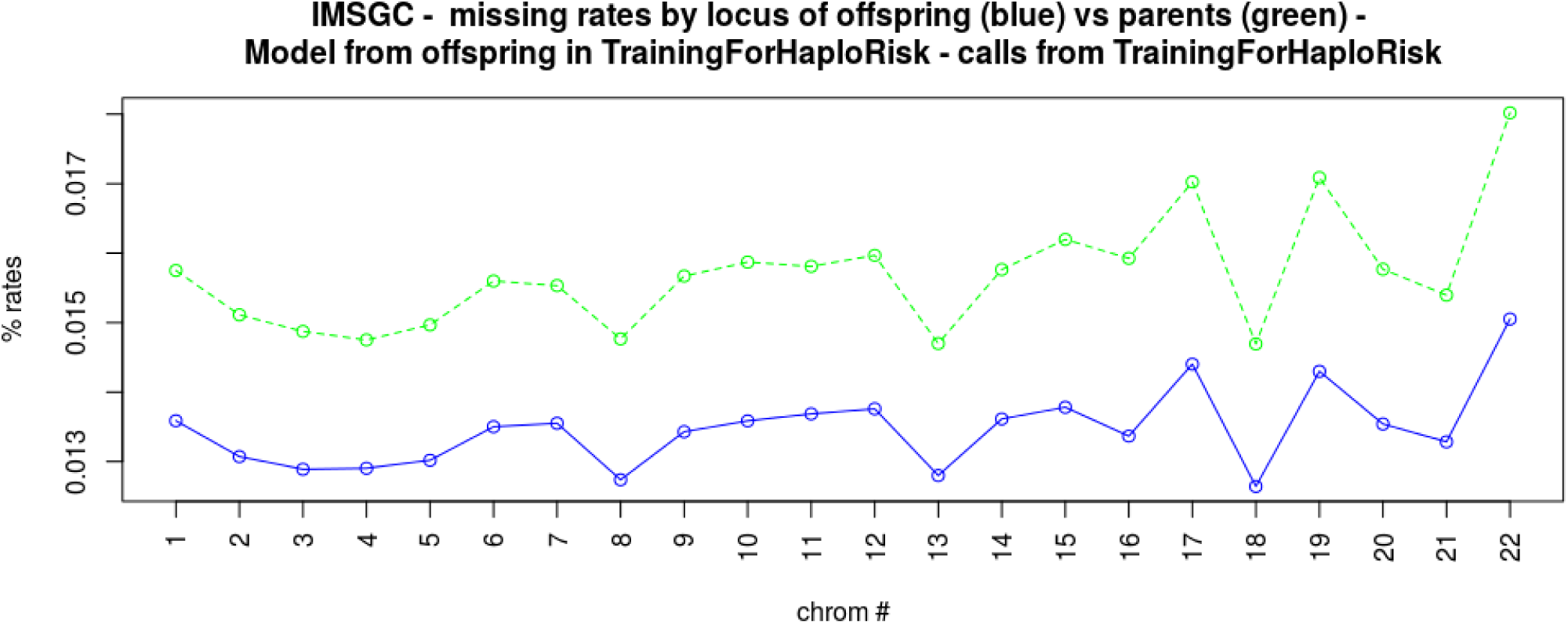
Missing rates in offspring (blue line) vs parents (green line) using only the offspring to learn the clustering model.

**Figure 19.**
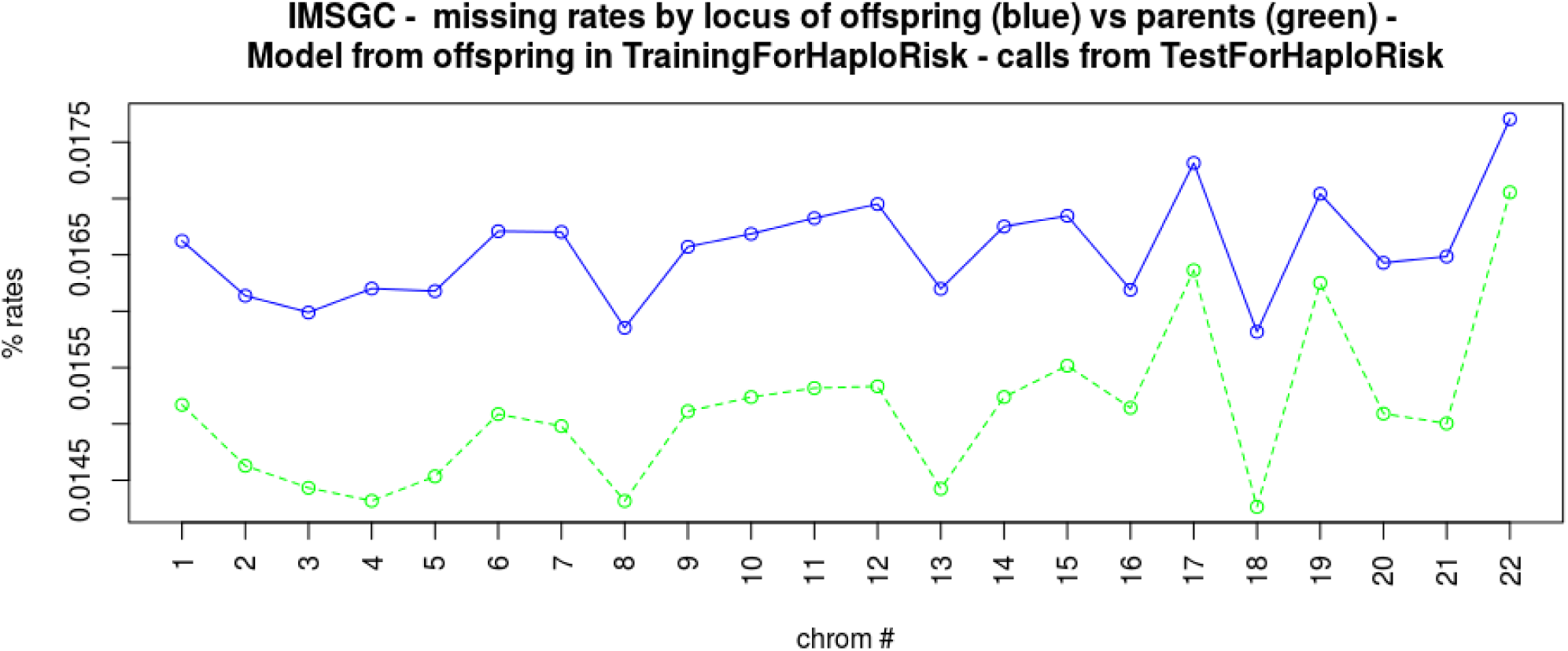
Missing rates in offspring (blue line) vs parents (green line) using an independent set of offspring data set to learn the clustering model.

We also tried other algorithm for clustering, different to the default Contrast Centers Stretchs (CCS), as we thought it could be better given what we knew so far about the missing calls. The algorithm just transformed intensities by their polar coordinates R vs θ (RvT). As a result, missing rates lowered in all the chromosomes and in both parents and offspring (data not shown). Figure 20 shows overall missing rates using CCS and RvT with the same data set to learn the clustering model and make the calls (Training), with independent sets (Test) and using all samples together for both clustering model learning and genotype calls. As it was expected, the lowest rates were reached when using all the samples for both tasks but this may be a reason for a bad generalization capacity of the predictor, i.e., a lower accuracy in a validation data set or in single trio families whether the model reached an exploitation phase for clinic purposes. When the same data set is used to learn the clustering model and make the calls (Training), the differential missing pattern between parents and offspring are inverted, I.e. rates are lower in offspring, regardless the clustering algorithm used. This result can be explained because of sample overfitting.

**Figure 20.**
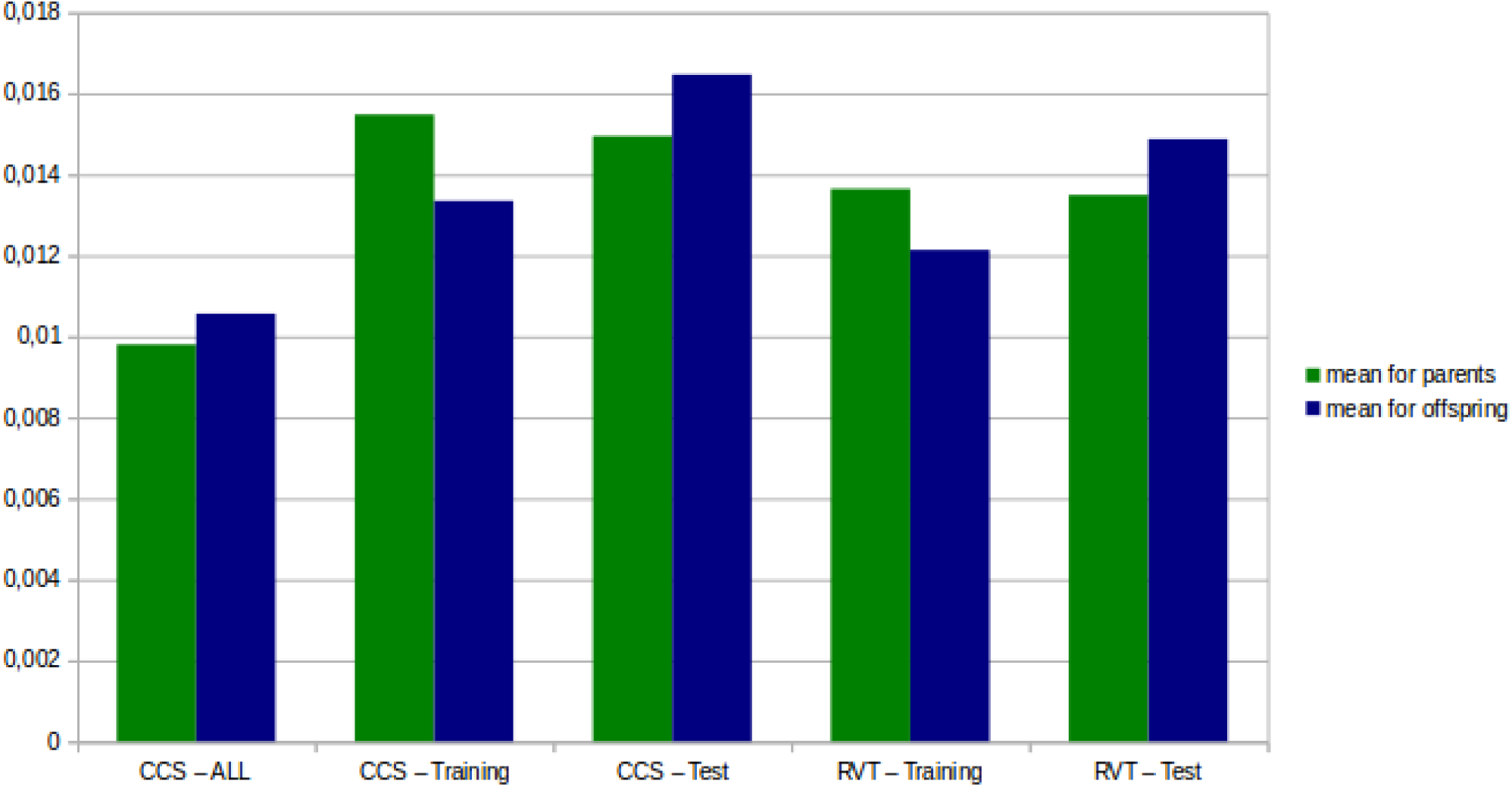
Missing rates in offspring (blue line) vs parents (green line) using only the offspring to learn the clustering model. The differential pattern of missing rates between parents and offspring is inverted, i.e., there are lower rates of missing genotypes in offspring, when the same data set is used to learn the clustering model and make genotype calls (“Training”), regardless the clustering algorithm used.

## 4. Discussion

At a first glance, missing rates returned by algorithms that perform genotype calls seem to be very low to have any impact on genetic predictors of complex diseases. However, this is against to our results and we need to understand why. Under the hypothesis of “Common Disease, CommonVariant” (CDCV) for complex diseases etiology, current missing rates are actually low. However, under the hypothesis of “Common Disease, Rare Variant” (CDRV), these rates may result very high. As it is believed that both hypothesis may coexist in the genetics of complex diseases, perhaps in different forms of a disease [25], we claim that the role that rare or vary rare variants play in complex diseases may explain the importance that missing rates have in the low performance of the current multiplex predictors.

Due to missing genotypes, and because genetic predictors for complex diseases should use genome-wide approaches to capture also the contribution of perhaps thousand variants with very small effect on the disease, the cT approach that we were applying by error to complete missing genotypes, may cause a low-risk variant of a small effect marker to become a high-risk variant of a larger effect marker, in such a way that we will have a trend to raise type I errors, with higher power and lower specificity. The same effect occurs with other approaches that imputes missing genotypes in order to increase transmissions of always the same allele. All these approaches are also imputing in a different way unaffected individuals (parents) and affected ones (offspring) and therefore can never being applied on a test data set, as a test data set has to be as much closer to a real situation for the predictor to me clinically usable. When we were aware of this highly biased way to complete missing genotypes we tried either to ignore missing or to complete in different ways but always at the same manner in parents and children but we had not success at all.

Therefore, if we do not specially consider missing genotypes or we force genotype calls to always make a call even if it is done by almost random guessing, detected risk variants may be biased towards the major allele, which is not a problem as far as it is also the high-risk variant. However, whenever a high-risk variant is associated with the mutant allele, as it is more difficultly ascertained and there are more missing calls, its result may be to underbias the effect that a variant has on the disease. Therefore, predictors built on data with missing genotypes or with algorithms that forced genotype calling towards the major allele, may have a trend to mostly type II errors, showing very low power and high specificity. This situation is in agreement with the state-of-the-art genomic profiling of complex diseases, in which AUC is much lower than it should be and it is caused by a lack of power [7]. A similar trend against the high risk allele whenever it is also the minor allele, may occur in algorithms used to perform missing imputation and haplotype reconstruction. As genetic causes of complex diseases may come from hundred thousands of variants with very small effects, we also wonder whether current procedures for QC may not only remove noise but also a true signal.

The singular distribution of intensities that were observed in three of the first five SNPs with more differences in missing rates between offspring and parents, is something to study in deep. Instead of using stringent QC procedures to remove Mendelian errors or to ignore missing genotypes, it may be worth to understand that raw intensities may not show the clear three-cluster patterns (one for each genotype), perhaps due to copy number variation, deletions or other different causes. If these different patterns of intensities are frequent and they are associated with the disease, we cannot ignore them if we pursuit to build accuracy predictors. Therefore, they should be first understood before studying the patterns of missing and their differences between affected and affected individuals.

## 5. Conclusions

Nowadays, more than a year since the second grant to support this work finished, we do not know whether to use trios to better reconstruct haplotypes, to collape SNPs using 2G as a way to enhance the information of each variable in a GRS [21], to compute risk for each genome-wide haplotype by a hwGRS and to combine them under the assumption of a recessive model [8] actually would outperform the predictive capacity of other state-of-the-art genetic predictors. All of them have shown to be much worse than what it would be expected according to disease prevalence and inheritance whose occurrence they try to predict [7]. However, there is an important result from all this work that we reached by pure chance when were imputing missing SNPs being unaware of that: rates of missing genotypes were so high (by considering the impact of many low-effect variants in complex diseases) that, if we defined the interval of all possible ways to impute them, as it was done for single SNP positions [22], we could move from models highly downbiased (very low sensitivity and very high specificity), to models highly upbiased (very high sensitivity at a cost of very low specificity). We have also shown that missing rates are, in all chromosomes, more frequent in offspring than in their parents in MS, asthma and autism, the three diseases among those that we used with a very high genetic component. This missing pattern was exactly the opposite one observed in a data set of control trios (CEPH from HapMap).

When the Human Genome Project finished on 2003, genetic profiling seemed to be an unquestionable consequence that would come very soon. However, we are right now far away to be able to build genome-wide genetic predictors of complex diseases that can be clinically usable [7]. The main cause usually given to explain this defeat, after so many years since the first human being was genotyped is the sample size. As in a complex disease there may be thousand variants with low effect and much more with very low effect, we would need larger data sets to capture this effect.

As a results of this work, we believe this generalized lack of success many not only be solved by increasing samples sizes. We believe missing genotypes follow an informative missing pattern related to the disease. The same seems to occur with Mendelian errors. We believe they are not random and we would need to look closer to these patterns in order to assess whenever a missing or wrongly called genotype is associated to the disease that is being analysed or it can be considered random by that disease. Only if we are able to identify differences between a missing at random pattern and an informative missing pattern to disease risk, we would be able to decide which and how missing genotypes should be imputed.

## Acknowledgments

We want to particularly acknowledge the patients with multiple sclerosis and the control subjects who provided the source for the MS-ES data set and to the Nodo Biobanco (Biobank) Hospitalario Virgen Macarena (Biobanco Sistema Sanitario Público de Andalucía) for its help and support in the gifts of clinical samples used in this work.

We thanks the IMSGC for providing to us the calling genotypes of their GWAS before we could access through dbGaP. We acknowledge to the University of California at San Francisco and specially to Prof Esteban González Burchard as principal investigator of the GALA I project, for giving to the University of Granada access to the family trios data set of genotypes (the Asthma data set in this work). We acknowledge Profs Paola Sebastiani from Boston University and Chris Gignoux from UCSF, for their help in this work. We also acknowledge the help of some colleagues at the University of Granada (V Potenciano, Prof S Moral, A Masegosa, S Torres-Sánchez and F Molina), to Profs Fuencisla Matesanz and A Alcina from IPBLN-CSIC, Granada and G Izquierdo-Ayuso and M García-Sánchez from Hospital Virgen Macarena, Sevilla, Spain and Prof. Mohamed Abd Allah Makhlouf from Suez Canal University.

